# Biofilm formation on glycated collagen modulates *Streptococcus mutans* bacterial extracellular vesicle production and cargo

**DOI:** 10.1101/2024.12.28.630307

**Authors:** Camila Leiva-Sabadini, Pablo Berríos, Paula Saavedra, Javiera Carrasco-Rojas, José Vicente González-Aramundiz, Mario Vera, Estefanía Tarifeño-Saldivia, Christina MAP Schuh, Sebastian Aguayo

**Affiliations:** Institute for Biological and Medical Engineering, Pontificia Universidad Católica de Chile, Santiago, Chile; Centro de Medicina Regenerativa, Facultad de Medicina, Clínica Alemana-Universidad del Desarrollo, Santiago, Chile; Departamento de Farmacia, Facultad de Química y de Farmacia, Pontificia Universidad Católica de Chile, Santiago 7820436, Chile; Departamento de Ingeniería de Minería, Escuela de Ingeniería, Pontificia Universidad Católica de Chile, Santiago, Chile; Department of Biochemistry and Molecular Biology, Faculty of Biological Sciences, University of Concepción, Concepción; School of Dentistry, Faculty of Medicine, Pontificia Universidad Católica de Chile, Santiago, Chile

## Abstract

*Streptococcus mutans* is the major microbial etiological agent of dental caries and can adhere to surfaces such as type-I collagen, present in dentin and periodontal tissues. Recent studies have characterized planktonic *S. mutans* bacterial extracellular vesicles (bEVs) and demonstrated environmental-induced changes due to sugar presence or pH alterations. However, to date there are no studies exploring if surface-derived changes - such as tissue glycation - can modulate bEV production in the context of oral biofilm formation in the elderly. Therefore, the aim of this work was to determine the role of biofilm formation and collagen glycation on the morphology and composition of *S. mutans* bEVs. For this, bEVs from *S. mutans* biofilms on native and glycated collagen surfaces were isolated, characterized, and compared to bEVs from planktonic cells. Nanoparticle tracking analysis and microscopy confirmed bEV production and showed that bEVs from biofilms are smaller in size and less abundant than those from planktonic cells. Furthermore, proteome analysis revealed that *S. mutans* biofilm formation on native and glycated collagen led to the enrichment of several key virulence proteins such as Eno, LuxS, Tpx, and ScrB. Also, a shift towards proteins involved in metabolic processes was found in bEVs following biofilm formation on collagen surfaces, whereas glucan metabolism proteins were overexpressed in vesicles from the planktonic state. These results demonstrate that biofilm formation, as well as the glycation of collagen associated with aging and hyperglycemia, can modulate bEV characteristics and cargo and could play a central role in *S. mutans* virulence and the development of diseases such as dental caries and periodontal disease.

## 1. Introduction

Dental caries is a widespread and persistent chronic health issue, affecting around 2 billion people worldwide. Dysbiosis in the oral microbiome favors the overgrowth of cariogenic microorganisms such as *Streptococcus mutans* leading to acid production and localized destruction of dental tissues^1,2^. Hence, dental caries is considered a biofilm-mediated disease, which undergoes various stages: it starts with the adhesion of initial bacterial colonizers and progresses to biofilm formation with the production of extracellular polymeric substances (EPS) and attachment of secondary colonizers^3^. Therefore, in recent years strong efforts are being made to understand how genetic and environmental factors such as diet, smoking or aging, among others, can modulate the behavior of oral biofilms in order to develop novel methods to prevent and treat the disease.

Among these factors, aging is of particular interest due to the significant and rapid growth of elderly populations worldwide. Furthermore, the aging process introduces distinct changes in the tissues of the oral cavity^4^ such as alterations in the collagen matrix of dentin through spontaneous non-enzymatic processes forming advanced glycation end-products (AGEs)^5^. These AGEs interact with dentinal collagen amino acids, particularly lysine and arginine, and alter the mechanobiological properties of the matrix^6–8^. Especially exposure to methylglyoxal (MGO) has been demonstrated to alter diameter, density, and the number of collagen crosslinks^9–12^. Most importantly, glycation reactions are intensified in the presence of hyperglycemia and oxidative stress, which are major factors in type-II-diabetes, smoking, and cellular aging^4,5^.

Given the crucial role of collagen in the structure of dentin and periodontal tissues, it is unsurprising that oral streptococci express several collagen binding proteins (CBPs) such as WapA and SpaP for *S. mutans*, and SrpA for *Streptococcus sanguinis*^13–16^. In this context, recent research suggests that age-related modifications of collagen (among other structural proteins) and glycation may contribute to early bacterial adhesion to oral tissues^17–19^. However, whether collagen glycation can alter other important virulence factors (e.g. biofilm formation) in *S. mutans* has not yet been explored. In recent years, extracellular vesicles (EVs) have gained considerable interest, especially in facilitating and orchestrating cellular communication, tissue organization and biofilm formation^20,21^. EVs are cell-derived structures with a size of 30 to 200 nm, containing diverse cargo including proteins, lipids, nucleic acids, and sugars^22,23^. In the oral cavity, both pathogenic and non-pathogenic microorganisms, are capable of releasing so-called bacterial EVs (bEVs)^24^. Their production and release have been shown to be influenced by environmental factors such as the presence of certain nutrients and specific sugars, hypoxia, or pH^25–27^. However, it remains unknown whether age-related changes in collagen – such as glycation - can influence the production and composition of bEVs from *S. mutans*.

Therefore, the aim of this work was to determine the role of collagen glycation on the morphology and composition of bEVs produced by *S. mutans* biofilms. We believe that understanding how biofilm formation and collagen glycation alter bEV production can shed light on their potential role in promoting dental caries in the elderly, paving the way for innovative prevention and treatment strategies in the future.

## 2. Methodology

### 2.1. Bacterial strains culture conditions

For all microbial assays, the well-characterized *S. mutans* UA 159 strain was employed. Stocks were kept at −80 °C and cultured on brain heart infusion (BHI) agar plates or culture medium at 37 °C under aerobic conditions.

### 2.2. Collagen-coating and surface glycation

To obtain a collagen coating, tissue culture plates were incubated with type-I collagen (rat tail, 3 mg/mL, Gibco) at a concentration of 0.5 mg/ml at 37°C for 60 min. Subsequently, the supernatants were removed and replaced by 1X phosphate-buffered saline (PBS) or glycated with 10 mM of methylglyoxal (MGO), as previously described^18^. Since glycation increases the autofluorescence of collagen, native and glycated surfaces were measured every 24 h for 4 days with a multimodal microplate reader (Synergy HT, Biotek), utilizing black 96-well plates (excitation/emission: 360 nm/460 nm). For the topographical characterization of the resulting collagen coatings, atomic force microscopy (AFM) imaging was carried out using SCOUT350 RAu (NuNano, UK, k=42 N/m) probes on a MFP 3D-SA AFM system (Asylum Research) in air, with a scanning velocity of 0.7 Hz and scan area of 10 x 10 μm with a resolution of 512 x 512 pixels.

### 2.3. Biofilm formation and growth on glycated collagen substrates

For all biofilm experiments, native and glycated type-I collagen coated 96-well plates were incubated for 72 h at 37°C. Wells were thoroughly washed with 1X PBS to remove any unreacted collagen or MGO molecules, and 50.000 CFU of *S. mutans* were inoculated into each well with 100 μL BHI for 24 h. Subsequently, the presence of biofilms on the collagen substrates was confirmed using a crystal violet (CV) staining. For this, supernatants were removed, and biofilms were washed with 1X PBS to remove loosely bound bacteria. Subsequently, plates were air dried at room temperature for 15 min, followed by drying in a 60°C oven for 30 min. Finally, plates were incubated with 0.1% CV for 15 min and subsequently washed with distilled water. Biofilms were eluded with 95% ethanol and absorbance was measured at 562 nm on a plate reader (Synergy HT, Biotek). Furthermore, *S. mutans* biofilm formation on collagen surfaces was confirmed with AFM using the above-described methodology.

### 2.4. Scanning electron microscopy (SEM) of bEV production by planktonic and biofilm-bound *S. mutans*

Planktonic bacteria and resuspended biofilm bacteria were fixed prior to SEM. All samples were centrifuged at 13.000 RPM (mySPIN 12, Thermo Scientific, US) for 5 min, and the resulting pellet was resuspended in 4 % paraformaldehyde for 40 min. Subsequently, the pellet was washed in 1X PBS and bacteria were transferred to a poly-L-lysine (PLL) coated cover slip. Samples were allowed to dry for 30 min, and dehydrated in a series of 25 %, 50 %, 75 % and 100 % ethanol solutions. Finally, coverslips were sputter coated with a 5 nm gold layer and imaged with a FEI Quanta FEG250 SEM.

### 2.5. bEV isolation from planktonic and collagen-bound *S. mutans* biofilms

For planktonic *S. mutans* bEV isolation, 10^7^ CFU/mL cells were incubated in 100 ml BHI for 24 h at 37 °C. Samples were then vortexed vigorously and centrifuged at 4.000 RPM (NUWIND NU-C-200R-E, NuAire) for 20 min at 4°C to sediment cells. The resulting supernatants were filtered using a 0.22 μm syringe filter. Subsequently, bEVs were isolated in two steps. First, the supernatant was concentrated with ultrafiltration (Amicon filtration system, Merck Millipore) at 4.000 RPM, 4°C, for 15 min. Then, bEVs were pelleted with ultracentrifugation (125.000 x g, at 4°C, 2 h), using a T-890 fixed angle rotor (Thermo Fisher Scientific). The resulting bEV pellets were resuspended in 1X PBS and stored at −80 °C until further experimentation.

For biofilm-derived bEV isolation, collagen-coated 12-well plates were inoculated with 2 mL of BHI medium and *S. mutans* at a cell density of 1×10^7^ CFU/mL for 37°C for 24 h. Following incubation, supernatants were removed together with detached bacteria, and the resulting biofilms were harvested with a cell scraper and collected in a Falcon tube with PBS 1X. For isolation of biofilm-derived bEVs, the method described above for planktonic bacteria was employed.

### 2.6. bEV morphological and microscale characterization

Following isolation, nanoparticle tracking analysis (NTA, NS300, Malvern Analytical) was performed to quantify vesicle concentration and size from 3 technical replicates across 3 independent bacterial isolations (n=3). A 1:20 dilution of each sample in particle-free PBS was measured in triplicate with a camera level of 12-14 for 20 s. Furthermore, bEV morphology was observed with transmission electron microscopy (TEM). Samples were transferred onto a copper grid, counterstained with 2% uranyl acetate for 1 min and dried at 60 °C for 20 min. Vesicles were visualized with a Talos F200C G2 system. Furthermore, analysis with AFM was performed utilizing an MFP-3D system (Asylum Research, US) as previously described^28^. Briefly, bEV samples were placed onto a 0.1 mg/mL PLL-coated cover slip and imaged with intermittent contact using SCOUT 350 RAu silicon AFM cantilevers with a spring constant of 42 N/m (NuNano, UK).

### 2.7. Protein quantification and proteomic analysis

Proteome analyses were conducted on three independent samples of planktonic bEVs and two independent samples derived from biofilm bEVs to elucidate compositional differences under planktonic and biofilm conditions. High-resolution liquid chromatography-tandem mass spectrometry (LC-MS/MS) was employed. Proteins were extracted, precipitated, and subsequently dissolved in 30 μL of 8 M urea and 25 mM ammonium bicarbonate. Reduction and alkylation were performed, followed by enzymatic digestion using trypsin (1:50 enzyme-to-protein ratio; Promega) at 37 °C for 16 h. Peptides were purified using disposable Evotip C18 columns (EVOSEP Biosystems) prior to LC-MS/MS analysis conducted on an Evosep One system coupled to a timsTOF PRO2 mass spectrometer (Bruker). Protein identification was carried out using MSFragger software (version 4.1), with proteomes from *Streptococcus mutans* UA 159 (UP000002512), *Homo sapiens* (UP000005640), and *Bos taurus* (UP000009136) retrieved from UniProt as the reference databases. Label-free quantification (LFQ) was performed using FragPipe-Analyst. MaxLFQ intensities were calculated following median-centered normalization and Perseus-type imputation. A global threshold of 20% for non-missing values and 60% per condition was applied to ensure robust identification of proteins across conditions, while allowing for specific variability conditions.

Differential expression analyses were conducted using the limma package for pairwise comparisons between conditions (Col vs. Plk, MGO vs. Col, and MGO vs. Plk). Proteins were considered differentially expressed if they met the criteria of an adjusted p-value < 0.1 and an absolute Log2 fold change > 1. Functional enrichment analyses were performed on upregulated and downregulated protein subsets using DAVID. The selected enrichment terms included GOTERM Biological Process (levels 3–5), GOTERM Cellular Component (levels 3–5), and UniProt Keywords (Molecular Function and Ligand categories). Data handling and visualizations were performed in Python.

## 3. Results and Discussion

### 3.1. Collagen-bound *S. mutans* biofilms produce EVs with specific morphological characteristics

To emulate age-associated glycation *in-vitro*, a previously published MGO incubation model was employed to modify collagen-coated surfaces^11,19^. AGE accumulation was monitored by collagen autofluorescence over a period of 96 h, demonstrating a significant increase in collagen glycation after exposure to MGO (**Figure 1A**). The maintenance of the collagen fibrillar structure following glycation was confirmed with AFM imaging. This revealed a well-conserved collagen matrix and characteristic D-banding periodicity due to its expected molecular staggering (**Figure 1B**). Previous work has confirmed similar observations following MGO incubation that are in line with our present findings^11^ and confirm that our glycation model is representative of the expected *in-vivo* changes in collagen while maintaining its biological morphology.

**Figure 1:**
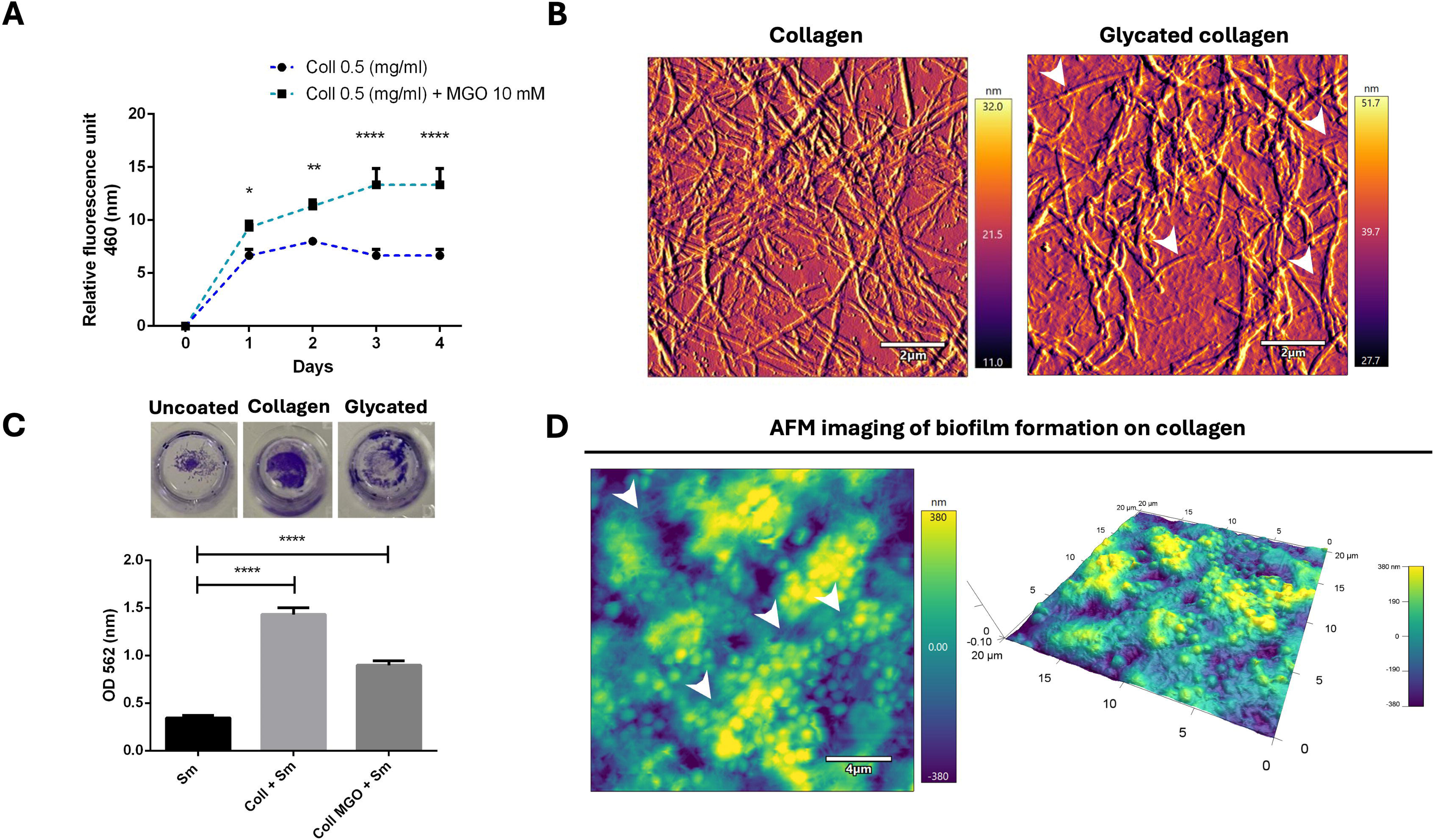
Methylglyoxal (MGO) glycation of type-I collagen coated surfaces allow *S. mutans* biofilm formation. (A) Autofluorescence quantification of glycation with 10mM MGO as a function of time (*p<0.05; t-test). (B) Representative AFM images of control and glycated collagen coatings showing the matrix network and maintenance of fibrillar structure and D-banding following glycation (white arrows). (C) Biofilm formation on uncoated, collagen coated, and glycated collagen coated wells (****p<0.001; ANOVA). (D) Height and 3D reconstruction images of a *S. mutans* UA 159 biofilm on collagen-coated substrates demonstrating the spacial interaction between bacterial cells and collagen fibers (arrows).

Biofilm formation on fibrillar collagen surfaces has been previously shown in studies with *Staphylococcus aureus*, and *Porphyromonas gingivalis,* among others^17,29^. Here we also demonstrate *S. mutans* biofilm formation on collagen-coated substrates (non-glycated and glycated) increases bacterial biomass compared to growth on uncoated plates (**Figure 1C**). This observation is expected considering that *S. mutans* express a range of collagen binding proteins that facilitate the colonization of collagen tissues^16^. Furthermore, AFM imaging confirms the formation of a *S. mutans* biofilm that is in direct physical contact with the collagen fibril matrix (**Figure 1D, arrows**), corroborating that the biofilm is effectively anchoring to the collagen surfaces and interacting with the organic matrix as expected.

Subsequently, the production of bEVs by planktonic and biofilm-bound *S. mutans* cells was assessed using SEM (**Figure 2**). Biofilm-derived cells were found to be embedded in EPS; nevertheless, the presence of bEVs on the surface of individualized *S. mutans* cells could be observed as previously shown^30,31^ (**Figures 2B** and **2C, arrows**). Similarly, planktonic cells were also seen to produce bEVs that were visible either on the surface of cells or expanding out of the bacterial cell wall (**Figures 2D** and **2E**). These observations confirm that bEV production is an active process within *S. mutans* growth and development, spanning from its growth as a planktonic cell to its surface attachment and adoption of a biofilm lifestyle. Following cell and biofilm growth, it was possible to isolate EVs from *S. mutans* from both collagen conditions by employing a combination of AMICON filtration and ultracentrifugation. These bacterial EVs were within the size range reported in previous literature^20,24,32^, with the largest size being found for planktonic EVs (PlkEVs), followed by bEVs from biofilms attached to collagen (ColEVs) and glycated collagen (MGOEVs) (124 nm, 112 nm, and 106 nm, respectively) (**Figures 3A** and **3B**). This is an interesting observation and comparable to the results reported by Johnston et al. that found larger EVs in *Pseudomonas aeruginosa* grown in planktonic state when compared to biofilm cells^33^. One could speculate that biofilm-derived bEVs are smaller due to the limited resources and spatial constraints of closely packed bacteria when compared to bacteria in suspension; thus, energy would be channeled towards biofilm maintenance rather than bEV synthesis^34^. Furthermore, planktonic bEVs were present in almost double the concentration than those isolated from biofilms; however, no differences were found regarding protein concentration across all three studied conditions (**Figure 3B**). It remains possible that the reduced yield observed for biofilm-derived vesicles is a result of bEVs being retained in the collagen matrix and not being liberated by mechanical preparation, similar to what has been recently described for matrix-bound vesicles^35,36^. The persistence of bEVs in the collagenous matrix after biofilm removal could have profound biological implications on maintaining chronic inflammation even after surface decontamination^37^ and should be further explored in the context of pathologies such as dental caries and periodontal disease.

**Figure 2:**
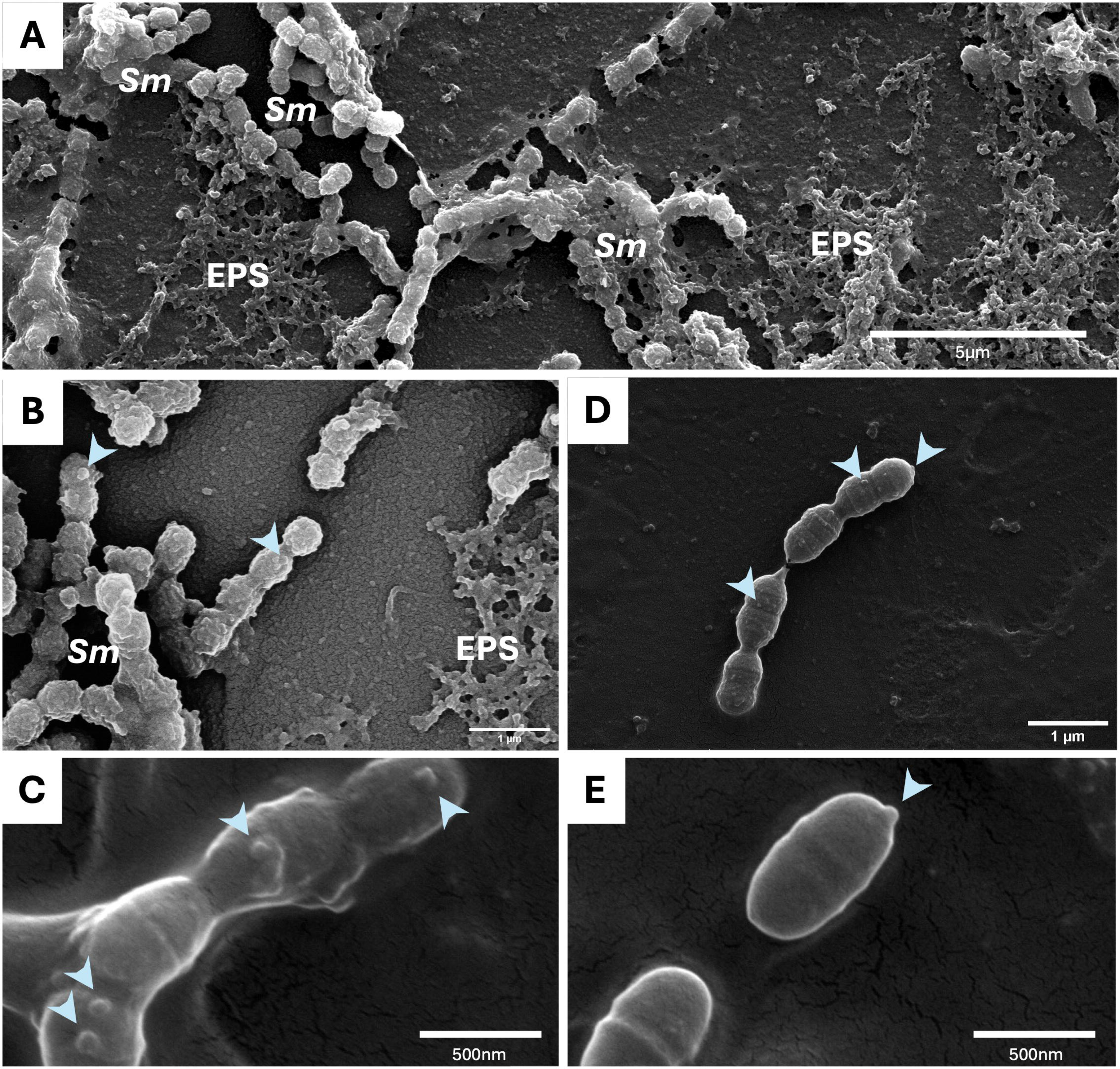
Ultrastructure of *S. mutans* extracellular vesicle (EV) production in planktonic and biofilm conditions. (A) Scanning electron microscopy (SEM) of adhered *S. mutans* biofilms showing extracellular polysaccharide (EPS) production and cellular aggregates. (B) and (C) High and low magnification of bEV production by biofilm-bound *S. mutans*, respectively. (D) and (E) High and low magnification of bEV production by planktonic (non-adhered) *S. mutans*, respectively.

**Figure 3:**
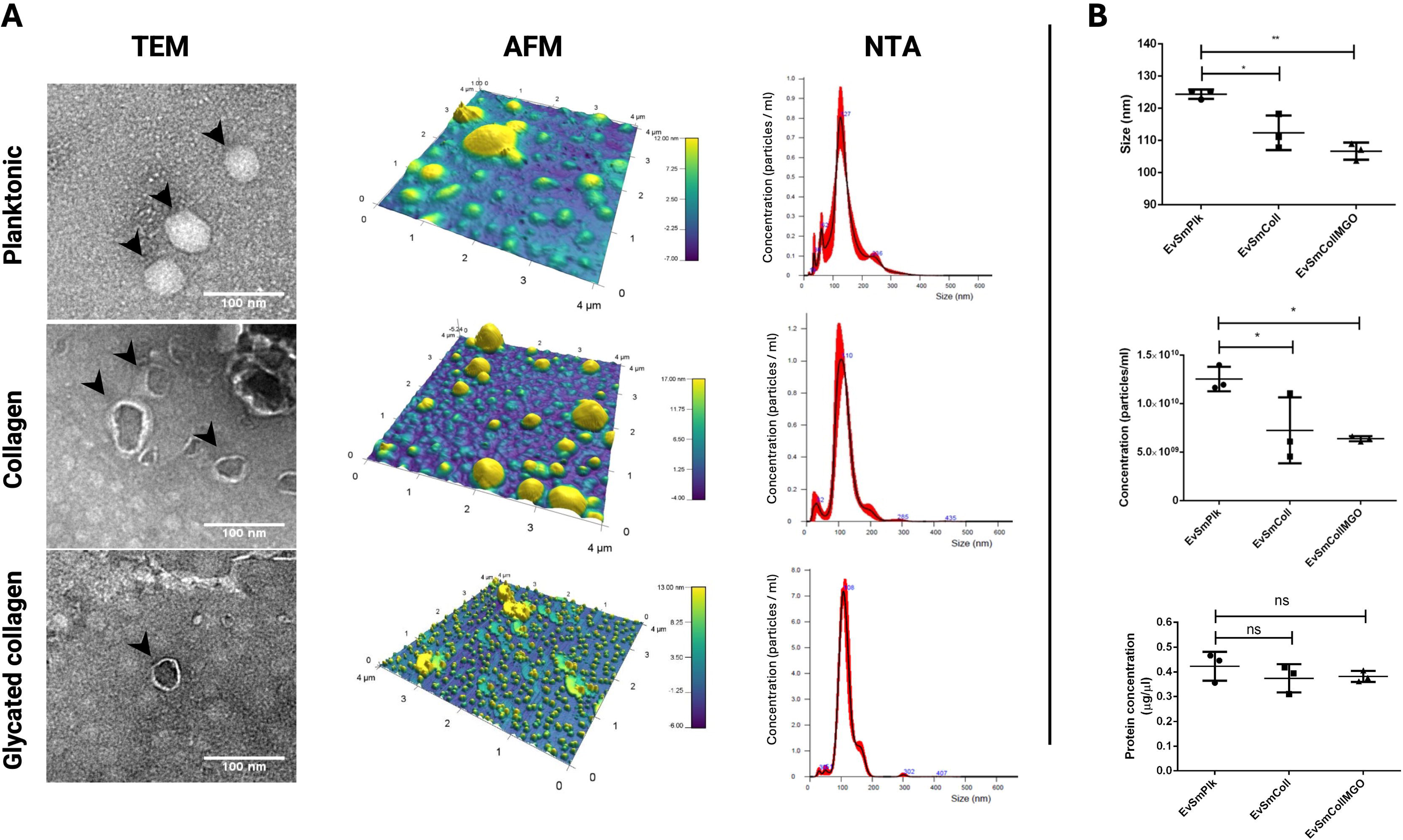
Morphological and microscopic characterization of *S. mutans* UA 159 bEV production on native and glycated type-I collagen substrates. (A) Transmission electron microscopy (TEM), atomic force microscopy (AFM), and nanoparticle tracking analysis (NTA) for bEVs isolated from each experimental condition. (B) Size distribution, particle concentration, and protein concentration for the investigated bEVs (*p<0.05, **p<0.01; ns: non-significant; ANOVA).

Furthermore, microscopic analysis with TEM confirmed the presence of bEVs in all the samples that were comparable to the morphology previously imaged bEVs from *S. mutans* and other microbial strains (**Figure 3E**, left arrows)^30,32,38^. Also, AFM imaging showed that bEVs from planktonic and native collagen-bound biofilms had a higher tendency to aggregate and cluster when deposited on the surface compared to MGOEVs (**Figure 3E**, right). This finding suggests substrate-specific changes in the surface lipid or protein composition of bEVs with the resulting vesicle physicochemical property changes^39^.

### 3.2. Proteome analysis of EVs demonstrates biofilm- and glycation-induced differences in EV cargo

Subsequently, a proteome analysis of EVs isolated from all three conditions was performed. Overall, a total of 247 proteins were found as cargo in PlkEVs, 457 in ColEVs, and 269 in MGOEVs. Among these, 201 proteins were found to be shared across all three groups (**Figure 4A**). The protein expression for frequently described *S. mutans* virulence factors such as GtfB, GtfC, Eno, LuxS, Tpx, and ScrB are illustrated in **Figure 4B**^40^. An interesting observation is that biofilm formation on collagenous surfaces triggers an increase in protein diversity within *S. mutans* EVs when compared to bacteria in suspension (**Figure 4C**). Furthermore, important changes in protein expression can be observed across the studied conditions and suggests that EVs from both biofilm conditions are more similar to each other than to EVs harvested from planktonic bacteria (**Figure 4C**).

**Figure 4:**
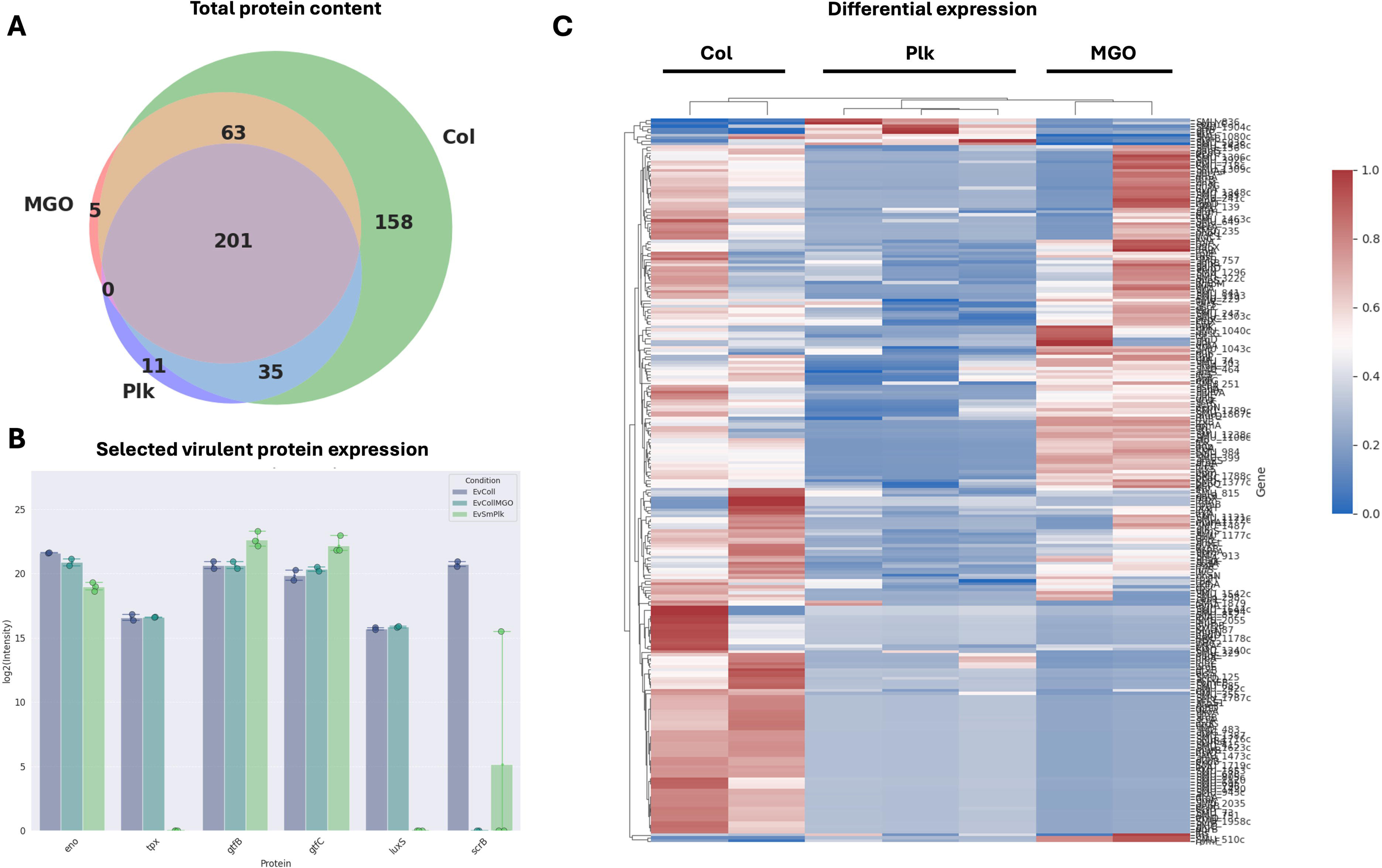
Proteomics *of S. mutans* bEVs demonstrate changes following biofilm formation on collagen and surface glycation. (A) Venn diagram illustrating the total number of proteins isolated in each sample. (B) Box plot illustrating the intensity for the relevant *S. mutans* virulence proteins GtfC, ScrB, GtfB, Eno, LuxS, and Tpx across the studied bEV groups. (C) Heat map of the overall protein expression from all 3 bEV conditions.

The differential expression analysis revealed 237 proteins differentially expressed between Plk and Col, 26 between Col and MGO, and 42 between MGO and Plk (**Figure 5A**, **Tables 1 and 2**). Thiol peroxidase (tpx), which catalyzes the reduction of hydrogen peroxide (H_2_O_2_) and is involved in stress oxidative tolerance^41^, was found to be overexpressed in collagen-bound biofilm bEVs (**Figura 5B**). This is clinically relevant in the context of cariogenic dysbiosis as it suggests that attachment to collagen can potentiate *S. mutans* inhibition of H_2_O_2_-producing commensal bacteria such as *S. sanguinis^42^* Click or tap here to enter text. and *Streptococcus gordonii* to promote its overgrowth on the dentinal surface. This is a crucial process for the formation of *S. mutans* microcolonies that can significantly reduce the local pH and promote demineralization of the tooth surface^43^. Furthermore, overexpression in bEVs of the molecular chaperones GroES and GroEL, involved in acid stress and heat shock^44,45^; thioredoxin reductase (TrxB), part of the thioredoxin system involved in reactive oxygen species (ROS) protection^46^; glutathione reductase (GshR), that also protects cells from ROS^47^; S-ribosylhomocysteine (LuxS) involved in EPS synthesis, biofilm formation, and quorum sensing^48,49^; and beta-ketoacyl-[acyl-carrier-protein] synthase III (FabH) related to fatty acid synthesis in bacteria^50^, was also found in both of the collagen-bound biofilm conditions. The vesicular overexpression of LuxS is quite relevant as it is known to be key in inter-species communication within the oral biofilm and has been shown to mediate the co-agreggation between *Porphyromonas gingivalis* and oral streptococci and induce periodontal ligament fibroblast inflammation^51^. Other biologically relevant proteins of interest that were overexpressed in bEVs isolated from biofilms include the signal recognition particle protein (Ffh) related to acid stress tolerance^52,53^, and the Clp-like ATP-dependent protease, ATP-binding subunit (Clp) that is known to have a protective role in the maintenance of the bacterial proteome^54^. Therefore, it remains possible that *S. mutans* bEVs are playing a role in modifying the local ecology towards dysbiosis following its attachment to native and glycated collagen in tissues, implying a role not only in dental caries but also periodontal disease modulation.

**Figure 5:**
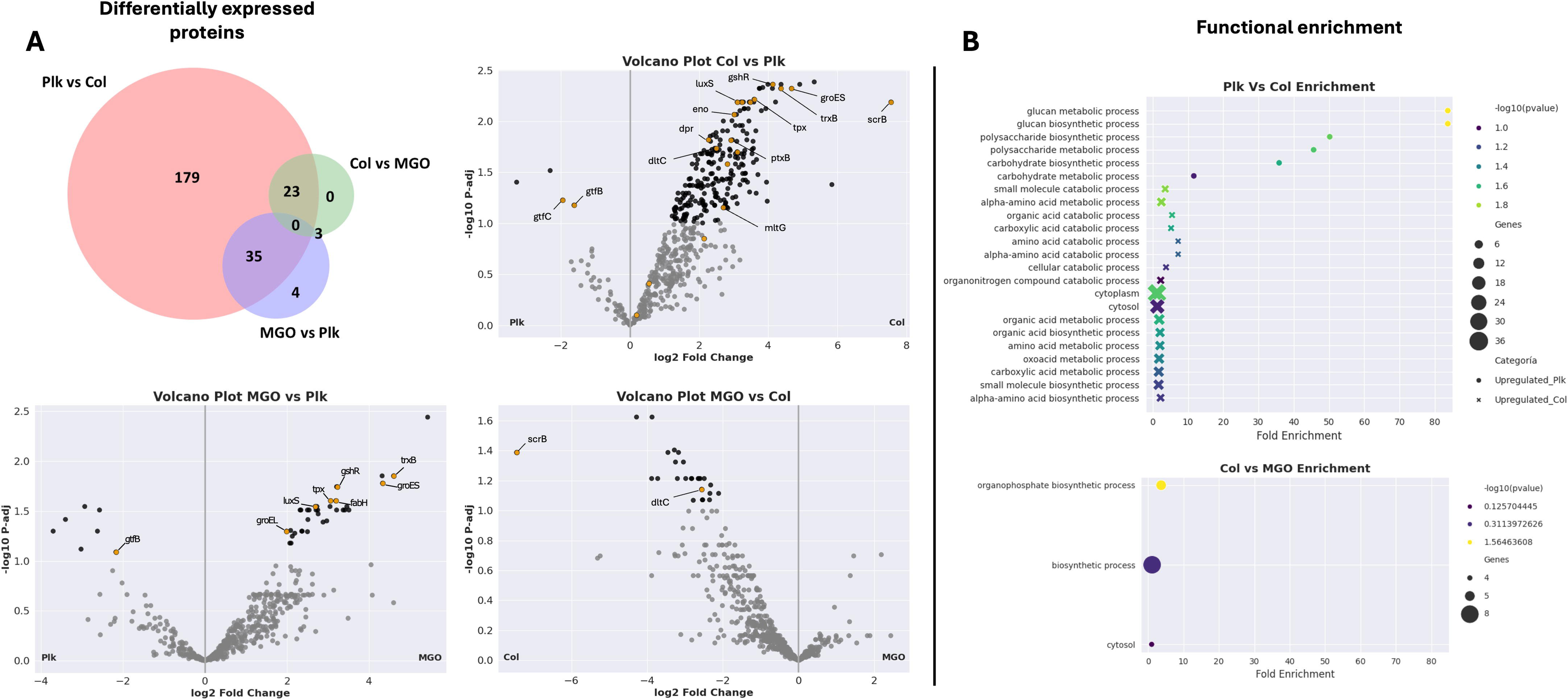
Differential expression of bEV proteins as a result of biofilm formation on native and glycated collagen substrates. (A) Venn diagram of differentially expressed proteins in Plk vs Col, MGO vs Plk, and Col vs MGO bEVs. The volcano plots illustrate the upregulation of proteins in bEVs from biofilms on glycated collagen compared to the planktonic and native collagen conditions. (C) Functional enrichment maps showing the main changes in bEV protein expression following collagen attachment and collagen glycation.

**Table 1:** Relevant overexpressed proteins in ColEVs and ColMGOEVs compared to PlkEVs.

| Protein ID | Gene | Protein description |
| --- | --- | --- |
| Q8DUQ4 | rpmA | Large ribosomal subunit protein bL27 |
| Q8DWB2 | pnp | Polyribonucleotide nucleotidyltransferase |
| Q8CWW5 | groES | Co-chaperonin GroES |
| Q8DVL7 | trxB | Thioredoxin reductase |
| Q8DUE5 | dapA | 4-hydroxy-tetrahydrodipicolinate synthase |
| Q8DUR5 | gshR | Glutathione reductase |
| P96995 | galE | UDP-glucose 4-epimerase |
| P95787 | atpA | ATP synthase subunit alpha |
| Q8DS44 | tyrS | Tyrosine--tRNA ligase |
| Q8DW30 | nifS | Cysteine desulfurase |
| Q8DUI4 | thyA | Thymidylate synthase |
| Q8DUK3 | tpx | Thiol peroxidase |
| Q8DTV8 | ung | Uracil-DNA glycosylase |
| Q8DSN2 | fabH | Beta-ketoacyl-[acyl-carrier-protein] synthase III |
| Q8DUA3 | SMU_1040c | Oxidoreductase, short-chain dehydrogenase/reductase |
| Q8DSP7 | greA | Transcription elongation factor GreA |
| Q8DSD8 | ssb | Single-stranded DNA-binding protein |
| Q8DSU4 | livF | Branched chain amino acid ABC transporter, ATP-binding protein |
| Q8DUH3 | clp | Clp-like ATP-dependent protease, ATP-binding subunit |
| Q8DSJ5 | SMU_1788c | Bacterocin transport accessory protein, Bta |
| Q8CWW6 | groEL | Chaperonin GroEL |
| Q54431 | ffh | Signal recognition particle protein |
| Q8DVK8 | luxS | S-ribosylhomocysteine lyase |
| Q8DS82 | nusG | Transcription termination/antitermination protein NusG |
| Q8DTX4 | SMU_1193 | Transcriptional regulator |
| I6L8Y7 | treA | Alpha, alpha-phosphotrehalase |
| Q8DW46 | SMU_229 | DhaL domain-containing protein |
| Q8DUE8 | SMU_984 | Peptidase C51 domain-containing protein |
| Q8DU47 | SMU_1106c | Phosphoglycerate mutase |
| Q8DVS0 | SMU_399 | C3-degrading proteinase |
| Q9X670 | pgi | Glucose-6-phosphate isomerase |
| Q8DTT5 | SMU_1238c | Uracil-DNA glycosylase-like domain-containing protein |
| Q8DSX9 | rplK | Large ribosomal subunit protein uL11 |

**Table 2:**
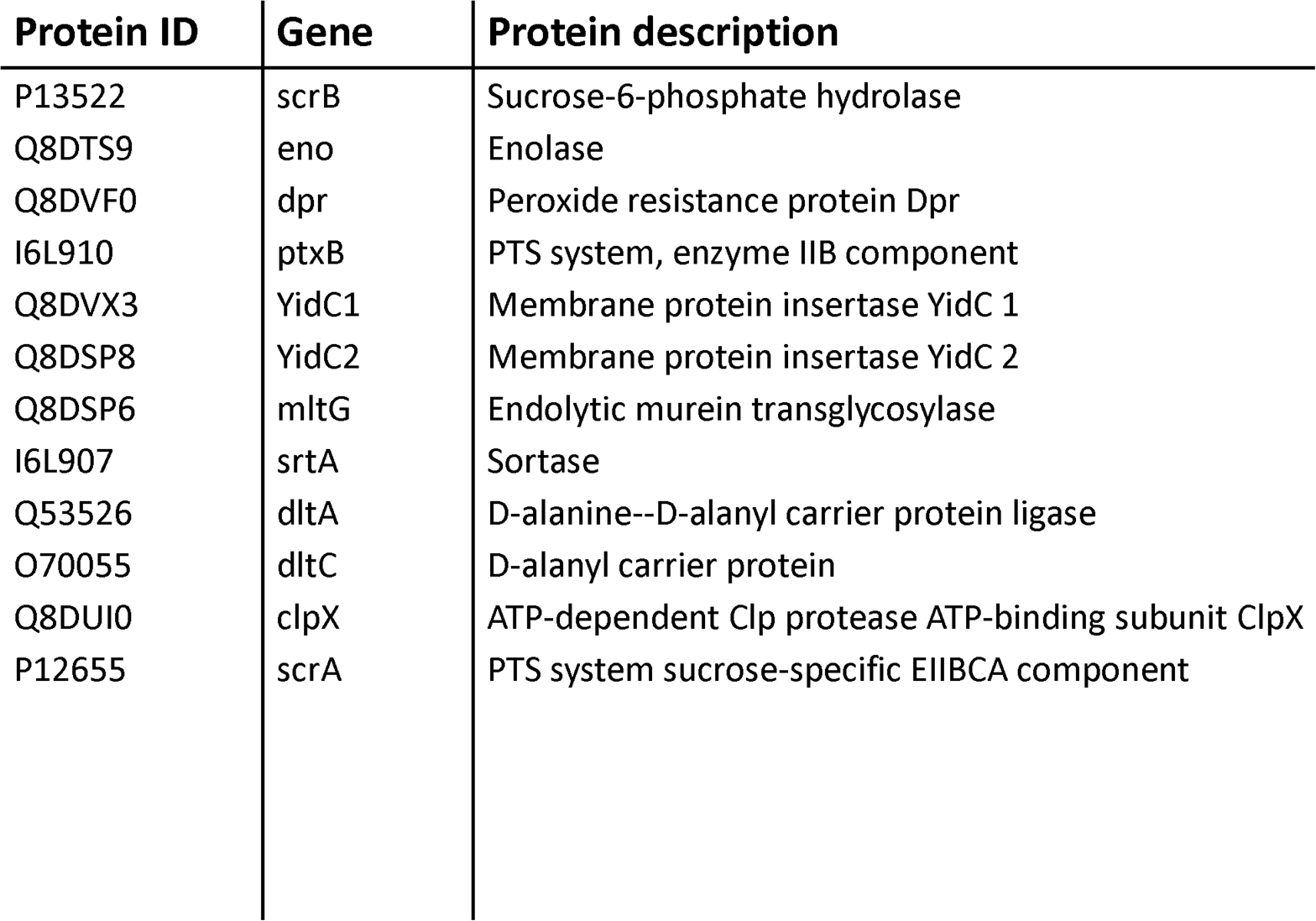
Selected overexpressed virulent proteins in bEVs from ColEV compared to PlkEVs.

| Protein ID | Gene | Protein description |
| --- | --- | --- |
| P13522 | scrB | Sucrose-6-phosphate hydrolase |
| Q8DTS9 | eno | Enolase |
| Q8DVF0 | dpr | Peroxide resistance protein Dpr |
| I6L910 | ptxB | PTS system, enzyme IIB component |
| Q8DVX3 | YidC1 | Membrane protein insertase YidC 1 |
| Q8DSP8 | YidC2 | Membrane protein insertase YidC 2 |
| Q8DSP6 | mltG | Endolytic murein transglycosylase |
| I6L907 | srtA | Sortase |
| Q53526 | dltA | D-alanine--D-alanyl carrier protein ligase |
| O70055 | dltC | D-alanyl carrier protein |
| Q8DUI0 | clpX | ATP-dependent Clp protease ATP-binding subunit ClpX |
| P12655 | scrA | PTS system sucrose-specific EIIBCA component |

In addition to those mentioned above, ColEVs displayed other relevant overexpressed proteins like enolase (Eno), a glycolytic enzyme that can translocate to the cell wall to act as a plasminogen receptor to bind human plasminogen^55,56^, and thus is believed to be involved in *S. mutans* remote tissue invasion and infective endocarditis^57^. Also, the increase of peroxide resistance protein (Dpr) involved in ROS protection^41^; the phosphor transferase system enzyme IIB component (PtxB) involved in biofilm development and acid response^58^; the ATP-dependent Clp protease ATP-binding subunit (ClpX) that modulates *S. mutans* virulence^59,60^; the sucrose-6-phosphate hydrolase (ScrB) and PTS system sucrose-specific EIIBCA component (ScrA), essential for sucrose internalization^61^; the membrane protein insertases (YidC1) and (YidC2) related to biofilm formation, protein secretion, and cell surface biogenesis^62^; and endolytic murein transglycolase (MltG) related to peripheral peptidoglycan synthesis^63^ were also found in bEVs isolated from biofilms attached to native collagen surfaces. Furthermore, bEVs from this condition displayed an overexpression of sortase (SrtA), a highly relevant enzyme that enables the anchoring of collagen-binding proteins such as SpaP and WapA to the *S. mutans* cell wall, and as such, can promote *S. mutans* adhesion and subsequent biofilm formation on collagen surfaces^16,64,65^. Regarding the comparison between the non-glycated and glycated collagen biofilm groups, an overexpression of ScrB and of the d-alanyl carrier protein (Dltc) associated with lipoteichoic acid metabolism^49,66^ was also observed (**Figure 4C**).

### 3.3. Biological relevance of *S. mutans* bEV modulation as a function of surface attachment and glycation

As a final step, an enrichment analysis of the *S. mutans* bEV proteome as a function of biofilm formation was carried out (**Figure 5B**). It was found that proteins involved in glucan synthesis processes were enriched in bEVs obtained from planktonic conditions compared to bEVs obtained from biofilms. In addition, biofilm formation on collagen promoted the packaging of proteins involved in several metabolic pathways within bEVs. Finally, collagen glycation was found to reduce the enrichment of biosynthetic proteins inside bEVs compared to the ones obtained on native collagen surfaces (**Figure 5B**). These findings suggest that when *S. mutans* bacteria are in a planktonic state – such as floating in saliva or crevicular fluid –they overexpress Gtfs in their bEVs to promote EPS formation in their surrounding environment with the goal of attaching to surfaces and forming a biofilm^40^. Once adhered, however, they switch to packaging metabolic proteins in order to kickstart biofilm formation and quorum sensing by the surrounding bacterial cells. This in conjunction with a higher content of Tpx can lead to a competitive advantage for *S. mutans* versus other early-colonizing commensal oral streptococci^42^. From our current results, it seems that bacterial adhesion alone is enough to induce changes in the secretory profile of *S. mutans* bEVs without the need of other environmental changes such as nutrient availability, flow, or pH alterations. It is well known that bacterial adhesion to surfaces and biofilm formation cause relevant changes in microbial transcription compared to planktonic cells^67^. Our present work demonstrates that these changes also hold true regarding the packaging of proteins in bEVs by *S. mutans*. To the best of our knowledge, this work is the first to show that biofilm formation on collagen can modulate the protein content of *S. mutans* EVs and suggests that this process is an important virulence factor in the development of oral diseases.

On the other hand, saliva poses a significant challenge to bacterial communication and survival due to the presence of redox-active molecules, proteases, and immune components^68,69^. In this context, proteolitic enzymes can degrade bacterial proteins involved in bacterial communication and biofilm formation, and secretory immunological molecules can rapidly neutralize bacterial surface and secreted proteins. These environmental pressures likely drive oral bacteria into employing protective strategies such as packaging key proteins and molecules in bEVs to shield them from salivary inactivation. Therefore, it is no surprise that *S. mutans* bEVs contain many virulent proteins including LuxS, Gtfs, and TrxB, among others, which are probably being trafficked as part of quorum sensing mechanisms among bacteria. In this context, future work should explore if these bEVs also mediate inter-species communication and quorum sensing within polymicrobial biofilms and drive the establishment and progression of local and systemic infection. Furthermore, how these alterations in bEV cargo may act synergically with other glycation-induced changes that directly affect bacterial cells and biofilms – such as increased adhesion and extracellular DNA secretion – remains to be elucidated^18,70^.

## 4. Conclusion

*S. mutans* bEV production is modulated by both collagen attachment and surface glycation by MGO. Biofilm cells were found to produce smaller bEVs as well as reduced particle yields when compared to planktonic *S. mutans*, although biofilms on glycated surfaces resulted in less bEV aggregation. Furthermore, important changes in bEV protein cargo and expression were observed as a result of biofilm formation and glycation, including crucial virulence factors such as GtfB, GtfC, Eno, LuxS, Tpx, and ScrB that are involved in key processes associated with biofilm formation. Also, a shift towards the packaging of proteins involved in metabolic processes was found following biofilm formation on collagen surfaces. Overall, the present results suggest that biofilm formation on both native and glycated collagen surfaces modulate bEV production and cargo and may play an important role in *S. mutans* virulence and the development of oral disease.

## Acknowledgements

This work was supported by the ANID FONDECYT #1220804 and #1220803 grants. Camila Leiva-Sabadini is supported by the Beca ANID PhD Scholarship #21220799. We also thank the MELISA Institute for support with the proteomics work and the Advanced Microscopy Unit (UMA) UC for their assistance with bEV transmission electron microscopy.

## 5. Conflict of interest statement

The authors have no conflicts of interest to report for this work.

## Notes

### Competing Interest Statement

The authors have declared no competing interest.

